# Distinct Computational and Temporal Mechanisms Underlie the Joint Effects of Motivation and Working Memory on Perceptual Sensitivity

**DOI:** 10.64898/2026.07.10.737658

**Authors:** Gargi Bhattacharjee, Shilpa Dang

## Abstract

Human perception is continuously shaped by internal cognitive states, yet how motivation and working memory jointly influence perceptual sensitivity remains poorly understood. Here, we combined behavioural experiments, computational modelling, and pupillometry to determine whether motivation enhances perception by amplifying working memory (WM)-driven facilitation or whether both exert independent influences. Participants performed a near-threshold visuospatial discrimination task under systematically manipulated motivational and WM states. Behaviourally, both motivation and WM independently improved perceptual performance, producing the greatest enhancement when both were present. A Bayesian generalized linear psychometric model revealed that these improvements were best explained by independent additive contributions to effective perceptual sensitivity, rather than motivational amplification of WM. Consistent with this computational framework, pupil dynamics tracked trial-by-trial fluctuations in effective perceptual sensitivity while revealing temporally dissociable influences of WM and motivation during perceptual decision making. Together, our findings demonstrate that the joint effects of motivation and working memory arise from distinct computational and temporal mechanisms, providing a unified framework for understanding top-down regulation of human perception.

## Introduction

On a busy street, while navigating through dense traffic, one can instantly detect a friend they are looking to meet. Likewise, a student scanning an exam sheet often becomes more precise in identifying relevant information under time pressure and motivation to perform well. These everyday scenarios illustrate that perception is not merely a passive registration of sensory input; rather, it is dynamically shaped by internal states such as working memory (WM) and motivation. When task-relevant information is actively maintained and aligned with goal-driven incentives, perceptual systems appear to operate at an enhanced level ^1,2^.

A substantial body of behavioural work shows that working memory biases visual attention toward stimuli that match maintained information ^3,4^, with effects emerging rapidly and potentially influencing early stages of visual processing ^5,6^. Consistent with this, improved discrimination accuracy has been reported when target features correspond to WM content ^7–9^, often interpreted as reflecting enhanced perceptual sensitivity to target features. However, these benefits were typically observed in multi-item displays with distractors, raising the possibility that they reflected reduced noise rather than genuine enhancement of sensory signals ^7,10^. Addressing this limitation, more recent findings using single-item displays without distractors demonstrated that WM can still improve discrimination accuracy for target features (gap side) corresponding to memory-matching content (colours), even in the absence of stimulus competition ^11,12^, suggesting a direct influence of WM on perceptual representations. These effects have been shown not to be driven by passive priming—defined as residual facilitation arising from recent stimulus exposure—but instead to reflect active WM-driven facilitation, whereby task-relevant information is deliberately maintained to bias processing toward matching stimuli. It has been shown that WM-driven facilitation enhances perceptual processing and may manifest as increase in perceptual sensitivity, potentially operating through faster orienting and/or improved sensory representation of target features ^5,6,13,14^.

Together, these findings support the view that working memory can directly influence perceptual performance, while leaving open how such enhancements interact with other modulatory factors such as motivation. Complementing this line of work, research on motivational effects has shown that both reward and loss contexts can influence perceptual performance ^15,16^. In particular, increases in reward value have been associated with enhanced perceptual sensitivity, suggesting that motivation can modulate perceptual representations ^17,18^. To the best of our knowledge, only a single previous study has examined the combined effects of attention and motivation on visual task performance through systematic and independent manipulation of attentional and motivational influences ^19^. Critically, although independent research studies have shown that both working memory and motivation can enhance perceptual performance, their joint influence and the mechanisms through which they modulate perceptual processing remains insufficiently understood. One possibility is that motivation amplifies WM-driven facilitation ^20^, in turn leading to enhanced perceptual performance. Alternatively, motivation and WM may exert independent additive influences on perceptual performance, resulting in greater enhancement than either factor alone. Addressing this question requires experimental paradigms that isolate perceptual processing while systematically manipulating both motivational and working memory states.

In the present study, we investigated the joint effects of motivation and visuospatial working memory on visual perception using a gap discrimination task adapted from a prior WM-based paradigm ^11^, which examined the influence of visual WM content (e.g., red or blue colours). Critically, we introduced two key modifications. First, rather than manipulating feature-based WM content (e.g., colours), participants maintained spatially specific information in visuospatial WM. Spatial representations are fundamental to visual processing because they enable the visual system to organize, prioritize, and efficiently process behaviourally relevant information across the visual field ^21,22^. Thus, we focused on visuospatial WM, allowing us to test how location-based memory representations facilitate perceptual processing in the visual field. Second, we incorporated motivational manipulations through neutral, gain, and loss contexts, enabling us to test how motivational state modulates perceptual processing alongside WM-driven facilitation. Participants performed the gap discrimination task under systematically manipulated conditions: WM with motivation, motivation only, WM only, and a neutral baseline, allowing dissociation of their individual and joint contributions to perceptual discrimination performance.

To examine the mechanisms underlying the joint effects, we employed behavioural, computational, and physiological approaches. Behaviourally, perceptual performance was quantified using statistical measures, including discrimination accuracy, reaction time, and efficiency score. Computationally, a Bayesian generalized linear psychometric model ^23^ estimated trial-by-trial fluctuations in effective perceptual sensitivity associated with motivational modulation and WM-driven facilitation. In addition, pupil diameter was recorded as a physiological index of arousal and cognitive effort during perceptual processing ^24,25^. Together, this framework allowed us to test whether motivation amplifies WM-driven facilitation to enhance perceptual sensitivity or whether motivation and WM exert independent additive influences, thereby providing mechanistic insight into how these internal states jointly shape perception.

Understanding these mechanisms may provide insight into how human perceptual systems flexibly adapt to real-world demands ^26^ and progressively exceed baseline performance limits as cognitive and motivational factors align ^27^, while also informing perceptual dysfunctions associated with disorders involving impairments in cognition and motivation ^28,29^.

## Results

### Behavioural Results

Participants completed three task components during a single experimental session: a perceptual threshold task to estimate participant-specific threshold stimulus duration, followed by the main perceptual discrimination task under visuospatial working memory and motivational state manipulations (hereafter, main task), and interleaved memory probe trials within the main task to verify active maintenance of the retro-cued spatial location.

In the perceptual threshold task, participant-specific threshold presentation duration for the peripheral Landolt stimulus were estimated using a double random interleaved staircase procedure combined with psychometric fitting (see *Methods* and **Fig. S1**). On each trial, stimulus presentation duration was adaptively varied between 100-180 ms based on participant performance using two randomly interleaved staircases ^30^. Performance across durations was subsequently fitted using a sigmoidal psychometric function, from which the presentation duration corresponding to 70% discrimination accuracy was estimated for each participant. **Figure S1** illustrates the trial structure of the threshold task and a representative psychometric fit from a single participant demonstrating threshold estimation. Across participants, the mean threshold duration corresponding to 70% discrimination accuracy was 142 ± 4 ms (mean ± SEM), providing participant-specific perceptual threshold duration within the range for use in the main task.

In the main task, participants performed the experiment under four principal conditions: performance-contingent monetary incentives combined with WM {Motivation, Cued}, motivational incentives without WM facilitation {Motivation, Non-cued}, WM without incentives {Neutral, Cued}, and a baseline condition lacking both motivational incentives and WM facilitation {Neutral, Non-cued} (see *Methods* and **Fig. 1A, B**). This design enabled isolation of the independent and joint contributions of motivational state and working memory relative to the baseline condition lacking both factors. Each trial began with sequential peripheral cue presentations followed by a retro-cue indicating the spatial location (out of 16 peripheral locations) to be actively maintained in working memory. After a delay period, participants performed a rapid perceptual discrimination task in which a briefly presented peripheral Landolt stimulus (participant-specific threshold duration; range: 100-180 ms) required a left/right gap discrimination response, allowing assessment of perceptual processing under different motivational and WM states. Motivational states consisted of neutral, gain, and loss contexts, in which correct or incorrect responses were associated with no monetary consequence, monetary reward, or monetary penalty, respectively. In cued trials, the perceptual target (Landolt stimulus) appeared at the retro-cued, i.e. actively maintained WM location; whereas in non-cued trials, target presentations were either at uncued or mismatched locations. Uncued locations referred to previously presented but non-maintained spatial positions capturing passive priming effects ^11^, whereas mismatched locations referred to positions not presented during the cue sequence. Further task details are provided in *Methods*.

**Figure 1.**
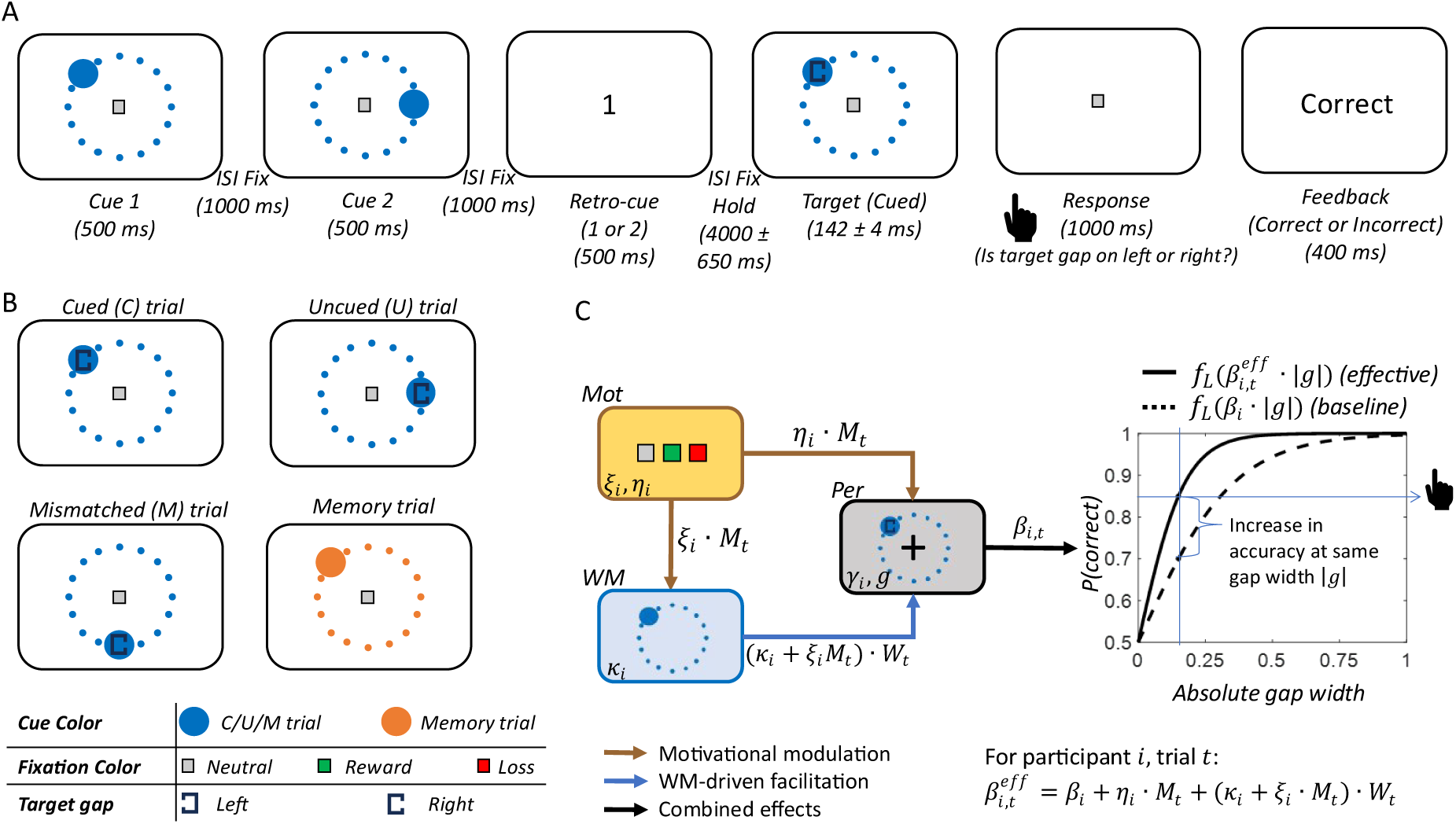
Experimental paradigm and computational framework. **(A)** *Trial structure of the main task.* Each trial began with the sequential presentation of two spatial cues (Cue 1 and Cue 2; 500 ms each), separated by fixation intervals (1000 ms). A retro-cue (1 or 2; 500 ms) instructed participants which location (Cue 1 or Cue 2, respectively) to maintain in working memory (WM). Following a delay period (4000 ± 650 ms), a target stimulus appeared at one of the peripheral locations (142 ± 4 ms). Participants indicated whether the gap in the perceptual target (a Landolt stimulus) was located on the left or right side within a 1000 ms response window. Feedback indicating whether the response was correct or incorrect was then presented for 400 ms. Trials were separated by a jittered inter-trial interval (2000 ± 460 ms). **(B)** *Working memory trial types and motivational conditions.* Main-task trials were indicated by blue-coloured cues. In cued (C) trials, the target appeared at the retro-cued location. In uncued (U) trials, the target appeared at one of the presented but non-cued location. In mismatched (M) trials, the target appeared at a location that had not been presented during the sequential cue period. Memory probe trials were interleaved throughout the experiment to assess working memory performance. These trials followed the same sequence as main task trials but omitted the perceptual target. Instead, participants judged whether a probe location matched the retro-cued location. The probe location could correspond to a cued, uncued, or mismatched location, and memory probe trials were indicated by orange-coloured cues. Motivational context was signalled by fixation colour: grey indicated neutral trials (no monetary outcome), green indicated gain trials (monetary reward), and red indicated loss trials (monetary penalty). **(C)** *Computational framework of perceptual decision behaviour.* Trial-wise motivational state (*M_t_*) and working-memory state (*W_t_*) jointly influence perceptual processing (gap discrimination) through distinct but interacting pathways. The motivational state exerts both a direct influence on perceptual sensitivity (*η_i_M_t_*) and an indirect influence through modulation of WM-driven facilitation (*ξ_i_M_t_W_t_*). The WM state selectively enhances processing of the target stimulus at the remembered spatial location (*κ_i_W_t_*). These influences combine with baseline perceptual sensitivity (*β_i_*) to determine effective perceptual sensitivity (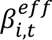) on each trial. Coefficients *β_i_*, *κ_i_*, *η_i_*, and *ξ_i_* denote participant-specific strengths of baseline perceptual sensitivity, WM-driven facilitation, direct motivational modulation, and indirect motivational modulation of WM-driven facilitation, respectively. Here, *i* denotes the participant index and *t* denotes the trial index. Effective perceptual sensitivity corresponds to the slope of the psychometric decision function (*f_L_*) and determines the probability of a correct response, *correct* , for a given level of sensory evidence (absolute gap width *g* ). The inset illustrates that, for a given level of sensory evidence, effective perceptual sensitivity yields greater perceptual accuracy than baseline perceptual sensitivity.

One participant was excluded due to poor perceptual discrimination performance in the { *eutral, on cued*} baseline condition (38.5% accuracy), falling below the predefined acceptable range (70% target threshold ± 20% tolerance). This resulted in a final sample of 31 participants for subsequent analyses. Within the main task, memory probe trials were randomly interleaved among perceptual discrimination trials to verify active maintenance of the retro-cued location. The memory probe trials followed the same sequence as the main task but omitted the perceptual target; instead, participants judged whether a probe location matched the retro-cued location. These trials were indicated using a distinct cue color from the main trials (**Fig. 1B**). Performance on memory probe trials was significantly above chance level across participants (mean ± SEM= 79.23% ± 2.04%), confirming reliable maintenance of the retro-cued spatial location throughout the task (*t*(30) 14.30, *p* 6.2 × 10^−15^, one-sample *t*-test against chance level of 50%).

To examine the independent and joint effects of motivational state and WM on perceptual performance in the main task, behavioural data were analysed using repeated-measures ANOVAs on efficiency score, accuracy, and reaction time (RT). Efficiency score, computed as accuracy/RT, was used as the primary behavioural metric because it jointly captures perceptual accuracy and response speed, thereby accounting for potential speed–accuracy trade-offs ^31^. No response trials were excluded from the analyses. A 3 × 3 repeated-measures ANOVA on efficiency score with factors Motivation (loss, gain, neutral) and WM location during target presentation (cued, uncued, mismatched) revealed significant main effects of Motivation (*F*(2,60) 43.61, *p* 2.02 × 10^-12^) and WM location (*F*(2,60) 28.18, *p* 2.35 × 10^-9^), (**Fig. 2A**). Critically, the Motivation × WM interaction was not significant (*F*(4,120) 0.81, *p* 0.52), suggesting largely additive influences of motivational state and WM on perceptual performance. Post hoc comparisons revealed significantly higher efficiency scores during motivational (loss, gain) relative to neutral conditions (*all p* < 0.01), whereas gain and loss conditions did not significantly differ from one another (*all p* > 0.55; **Table S1**). Similarly, cued trials produced significantly greater efficiency scores compared with both uncued and mismatched trials (*all p* < 0.01), whereas uncued and mismatched conditions did not differ significantly (*all p* > 0.10; **Table S1**). These findings indicated that motivational incentives and WM independently enhanced perceptual discrimination performance with no evidence for interaction effects. Because behavioural performance was statistically comparable between gain and loss conditions, as well as between uncued and mismatched conditions, subsequent analyses were performed using a reduced 2 × 2 design consisting of {Motivation, Cued}, {Motivation, Non-cued}, {Neutral, Cued}, and {Neutral, Non-cued} conditions. Gain and loss trials were collapsed into a single motivational condition, whereas uncued and mismatched trials were collapsed into a non-cued condition.

**Figure 2.**
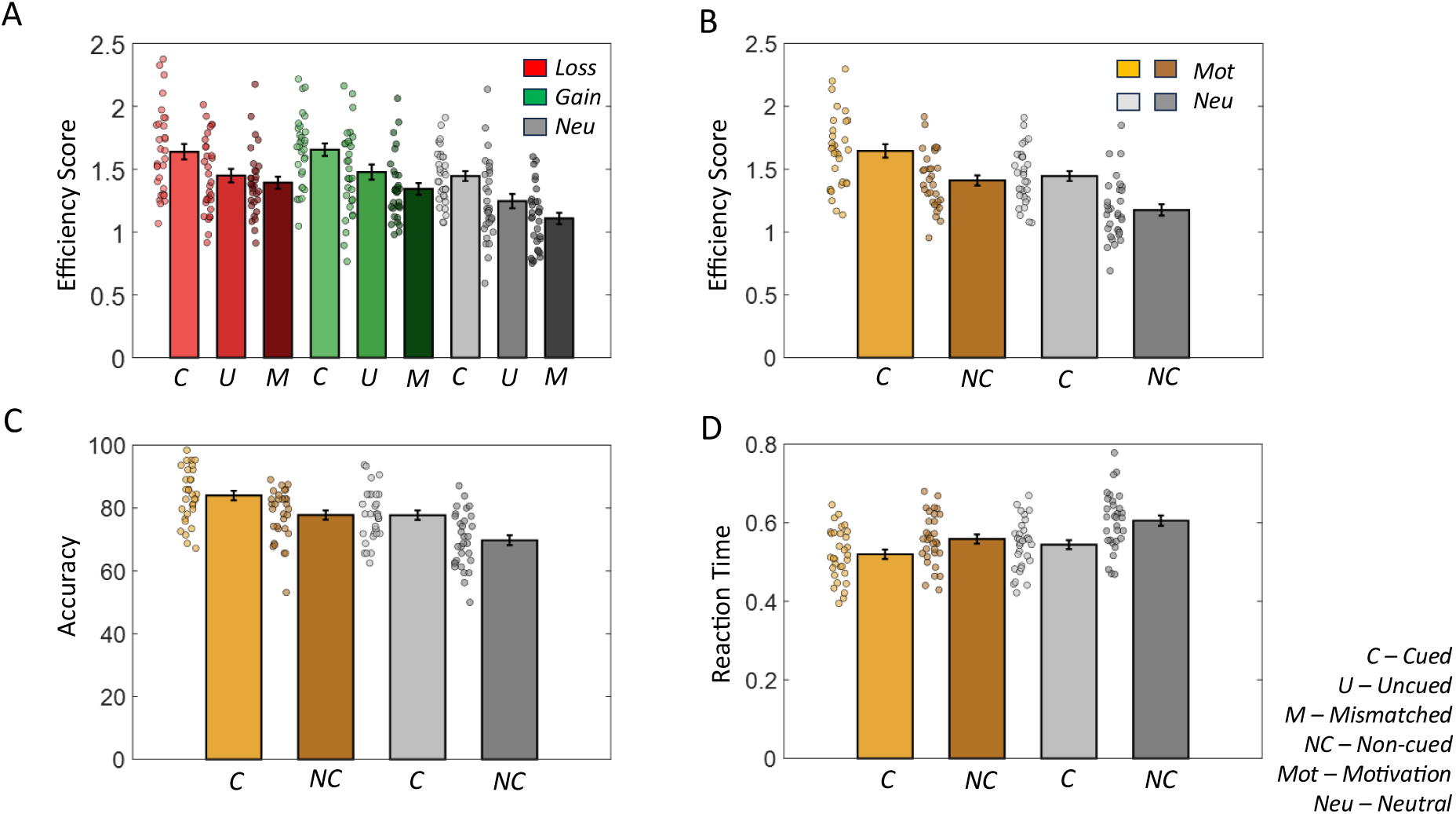
Behavioural effects of motivational state and visuospatial working memory on perceptual discrimination performance. (A) Mean efficiency score (accuracy/RT) across the full 3 × 3 experimental design comprising motivational state (loss, gain, neutral) and WM location during target presentation (cued, uncued, mismatched). (B) Mean efficiency score across the reduced 2 × 2 condition space used for subsequent computational modelling and pupillometry analyses. Gain and loss conditions were collapsed into a single motivational condition, whereas uncued and mismatched trials were collapsed into a non-cued condition. (C) Mean perceptual discrimination accuracy across the reduced 2 × 2 condition space. (D) Mean reaction times (RTs) for correct trials across the reduced 2 × 2 condition space. All data are shown as mean ± SEM. Markers represent individual participant data points *= 3* .

In the reduced 2 × 2 condition space, efficiency scores showed graded improvements across conditions, with both motivational state and WM independently enhancing perceptual performance relative to the {Neutral, Non-cued} baseline (**Fig. 2B**). Importantly, the combination of motivational state and WM ({Motivational, Cued}) produced the greatest behavioural enhancement, consistent with additive facilitative effects. A complementary 2 × 2 repeated-measures ANOVA similarly revealed significant main effects of Motivation (*F*(1,30) 68.66, *p* 3.00 × 10^-9^) and WM (*F*(1,30) 54.89, *p* 2.96 × 10^-8^), but no interaction (*F*(1,30) 0.66, *p* 0.42), confirming additive behavioural effects in the reduced condition space as well. (Detailed post-hoc statistics for the reduced 2 × 2 ANOVA are provided in **Table S2**.)

Accuracy analyses revealed a similar pattern (**Fig. 2C, Table S3**), with higher discrimination accuracy during motivational and cued conditions relative to neutral and non-cued conditions, respectively (main effects of Mot: *F*(1,30) 55.15, *p* 2.82 × 10^-8^, and WM: *F*(1,30) 23.61, *p* 3.4 × 10^-5^). No significant Motivation × WM interaction was observed (*F*(1,30) 0.56, *p* 0.46). RT analyses on correct trials demonstrated faster responses during motivational and cued conditions (main effects of Mot: *F*(1,30) 23.52, *p* 3.56 × 10^-5^, and WM: *F*(1,30) 70.99, *p* 2.11 × 10^-9^; **Fig. 2D**). A significant Motivation × WM interaction was additionally observed (*F*(1,30) 6.84, *p* 0.01), because WM-related effects (cued – non-cued) were larger in neutral than motivational trials as well as motivational effects (mot – neu) were larger in non-cued than cued trials (**Table S4**). The RT interaction indicated reduced scope for additional response-speed improvements when both WM and motivational facilitation were simultaneously present rather than either factor alone. However, the overall behavioural pattern remained broadly consistent with additive facilitative contributions of motivation and WM, particularly for efficiency score and accuracy measure.

### Computational Modelling

To characterize the computational mechanisms underlying perceptual decision behaviour during the main task, behavioural responses were analysed using a Bayesian generalized linear psychometric model ^23^. For each participant *i*, the model estimated trial-by-trial fluctuations in effective perceptual sensitivity (effPS; denoted by 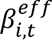 associated with motivational and WM states, thereby modulating gap discrimination accuracy (i.e., probability of correct responses) for the same level of sensory evidence (gap width) (**Fig. 1C**, see *Methods*). Specifically, effPS was modelled as a linear additive function of baseline perceptual sensitivity (*β_i_*), WM-driven facilitation (*κ_i_W_t_*), direct motivational modulation (*η_i_M_t_*), and indirect motivational modulation of WM-driven facilitation (*ξ_i_M_t_W_t_*). Trial-wise experimental conditions were represented using binary variables, with motivational state (*M_t_*) coded as 1 for motivational (loss/gain) trials and 0 for neutral trials, and WM state (*W_t_*) coded as 1 for cued trials and 0 for non-cued (uncued/mismatched) trials. The binary encoding of motivational and WM conditions was based on ANOVA results of behavioural efficiency scores, which revealed no significant differences between loss and gain conditions, as well as between uncued and mismatched conditions. The model therefore tested whether WM and motivation exerted independent additive influences on effPS, and/or whether motivation selectively amplified WM-driven facilitation.

A family of eight nested candidate models was constructed by systematically including or excluding direct motivational modulation of perceptual sensitivity (Mot→Per), WM-driven facilitation effects (WM→Per), and motivational modulation of WM-driven facilitation (Mot→WM) (**Fig. 3A**). Models were fitted using maximum a posteriori estimation and compared using random-effects Bayesian model comparison implemented in the Variational Bayesian Analysis (VBA) toolbox for MATLAB ^32,33^. Further details of model formulation, parameter estimation, and model comparison procedures are provided in the *Methods*.

**Figure 3.**
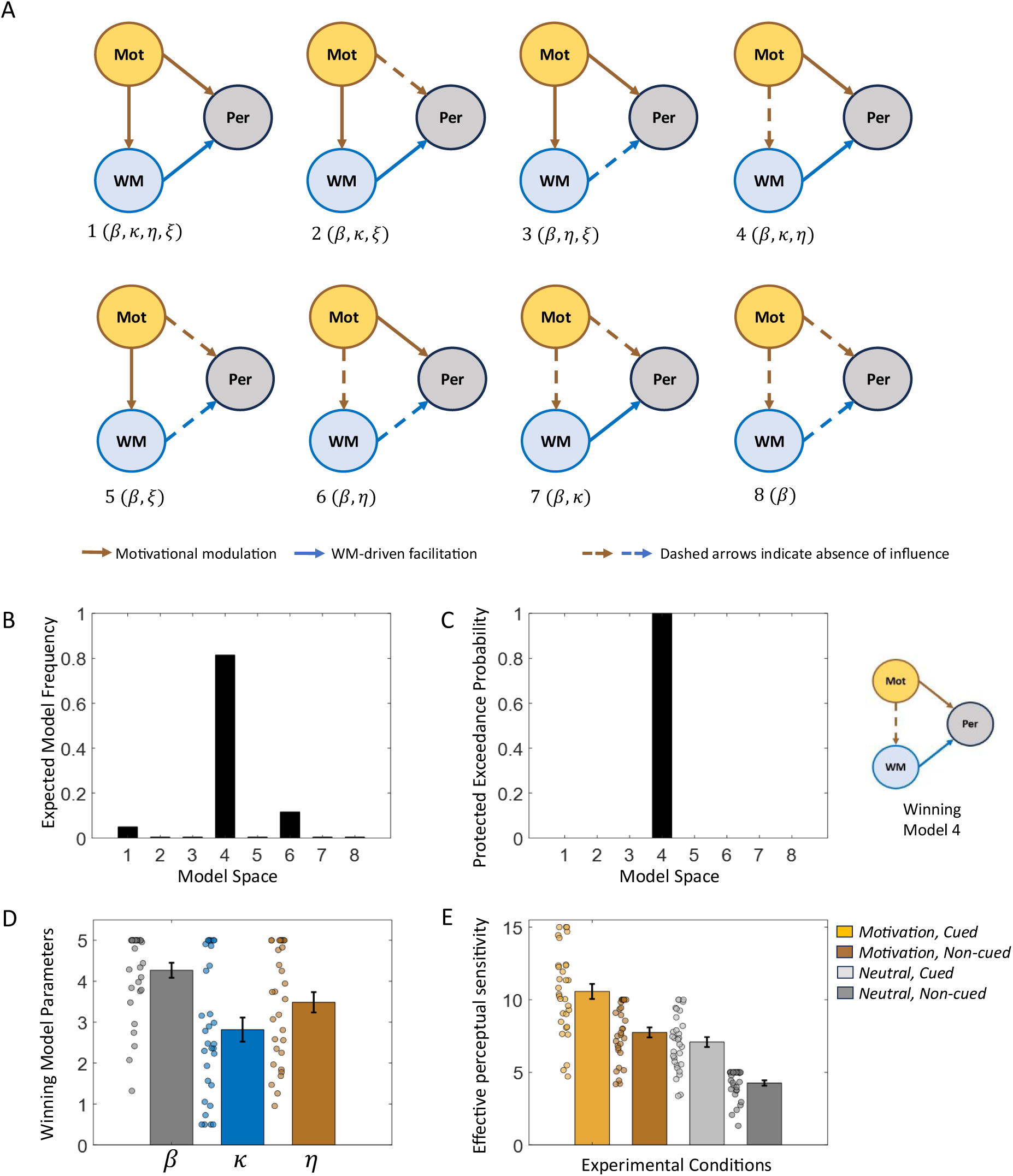
Computational modelling of perceptual decision behaviour under motivational and WM states. (A) Candidate model space comprising eight nested Bayesian generalized linear psychometric models. Models systematically varied the inclusion or exclusion of direct motivational modulation of perceptual sensitivity (Mot→Per; *η*), WM-driven facilitation effects (WM→Per; *κ*), and motivational modulation of WM-driven facilitation (Mot→WM; *ξ*), in addition to baseline perceptual sensitivity (*β*). Solid arrows indicate included influences, whereas dashed arrows indicate absent influences. (B) Expected model frequencies obtained from random-effects Bayesian model comparison across the model space. (C) Protected exceedance probabilities for all candidate models. The inset illustrates the winning model (Model 4), which included baseline perceptual sensitivity (*β*), WM-driven facilitation (*κ*), and direct motivational modulation of perceptual sensitivity (*η*), while excluding motivational modulation of WM facilitation (*ξ*). (D) Group-level parameter estimates for the winning model, including baseline perceptual sensitivity (*β*), WM-driven facilitation (*κ*), and motivational modulation of perceptual sensitivity (*η*). (E) Model-derived effective perceptual sensitivity across the reduced 2 × 2 condition space: {Motivation, Cued}, {Motivation, Non-cued}, {Neutral, Cued}, and {Neutral, Non-cued}. All data in (D) and (E) are shown as mean ± SEM. Markers represent individual participant data points ( *= 3* ).

Model comparison revealed that Model 4 (*β*, *κ*, *η*) provided the best account of behavioural performance across participants, exhibiting the highest expected model frequency (Ef = 0.81) and protected exceedance probability (pxp = 1.00) (**Fig. 3B, C**). The winning model included baseline perceptual sensitivity, WM-driven facilitation (Mot→Per), and direct motivational modulation of perceptual sensitivity (WM→Per), while excluding indirect motivational modulation of WM-driven facilitation (Mot→WM). These findings indicate that motivational states enhanced perceptual decisions primarily through a global increase in perceptual sensitivity rather than through selective amplification of WM-related facilitation. Instead, motivation and WM exerted independent additive influences on perceptual sensitivity, jointly contributing to enhanced discrimination performance. Group-averaged parameter estimates for the winning model were as follows (mean ± SEM): baseline perceptual sensitivity (*β*), 4.27 ± 0.18 (*t*(30) 23.29, *p* 0); WM-driven facilitation (*κ*), 2.82 ± 0.29 (*t*(30) 9.62, *p* 1.13 × 10^-10^); and motivational modulation of perceptual sensitivity (*η*), 3.48 ± 0.25 (*t*(30) 14.00, *p* 1.1 × 10^-14^) (**Fig. 3D**). Lapse rates were negligible (<10⁻⁷). All estimated parameters were positive, consistent with facilitative contributions of both WM and motivational state to perceptual sensitivity and in turn to discrimination performance.

Model-generated effective perceptual sensitivity estimates closely reproduced the empirical pattern of discrimination accuracy across motivational and WM conditions, indicating that the winning model captured the principal computational structure underlying perceptual decision behaviour (**Fig. 3E**). Specifically, **Figure 3E** illustrates a graded increase in effective perceptual sensitivity across conditions, demonstrating additive contributions of motivational state and WM-driven facilitation to perceptual discrimination performance. The {*Motivation*, *on cued*} and { *eutral*, *Cued*} conditions produced comparable increases in perceptual sensitivity individually; however, when both factors were simultaneously present, perceptual sensitivity increased substantially further, consistent with additive enhancement effects and supported by the absence of a significant Motivation × WM interaction (*F*(1,30) 0, *p* 1.00).

To assess model generalizability of the winning model, we performed leave-one-participant-out cross-validation (LOOCV). On each iteration, one participant was excluded, and group-average model parameters (*β*, *κ*, *η*) estimated from the remaining participants were used to predict the left-out participant’s trial-wise choice accuracy (correct vs. incorrect). This procedure was repeated for all 31 participants. Prediction accuracy was significantly above chance level (50%), indicating that the model generalized well to unseen participants (mean ± SEM = 78.51% ± 1.07%; *t*(30) 26.70, *p* 1.83 × 10^−22^).

To further validate the behavioural relevance of the computational framework, effPS estimates derived from the winning model were correlated with behavioural measures across 31 participants and 4 experimental conditions. effPS showed a strong positive correlation with discrimination accuracy (*r* 0.81, *p* < 0.001), indicating that increases in model-derived perceptual sensitivity were associated with improved behavioural performance. In addition, effPS was positively correlated with efficiency scores (*r* 0.75, *p* < 0.001)and negatively correlated with reaction times (*r* −0.43, *p* < 0.001), suggesting that greater model-derived perceptual sensitivity was associated with faster and more efficient perceptual decisions.

Overall, the computational modelling results suggest that motivational state and WM independently enhanced perceptual decision performance through additive increases in effective perceptual sensitivity. By modelling latent changes in perceptual sensitivity directly, this framework moves beyond descriptive behavioural measures and provides a mechanistic account of enhanced gap discrimination.

### Pupil dynamics track trial-by-trial fluctuations in effective perceptual sensitivity

We first examined whether pupil responses tracked trial-by-trial fluctuations in effective perceptual sensitivity (*β^eff^*), a latent computational variable integrating the influences of baseline perceptual sensitivity, working-memory facilitation, and motivational modulation. To this end, we applied a time-resolved general linear model (GLM-1) to z-scored pupil responses during the perception window (see *Methods*; **Fig. 4A, B, E**). The GLM-1 included a parametric effPS regressor and a general event-related (GER) regressor capturing pupil responses associated with task events common across trials. **Figure 4B** shows representative trial-wise raw values of regressor effPS for a single participant. This analysis allowed us to determine whether fluctuations in model-derived perceptual sensitivity were reflected in pupil dynamics independently of event-related responses.

**Figure 4.**
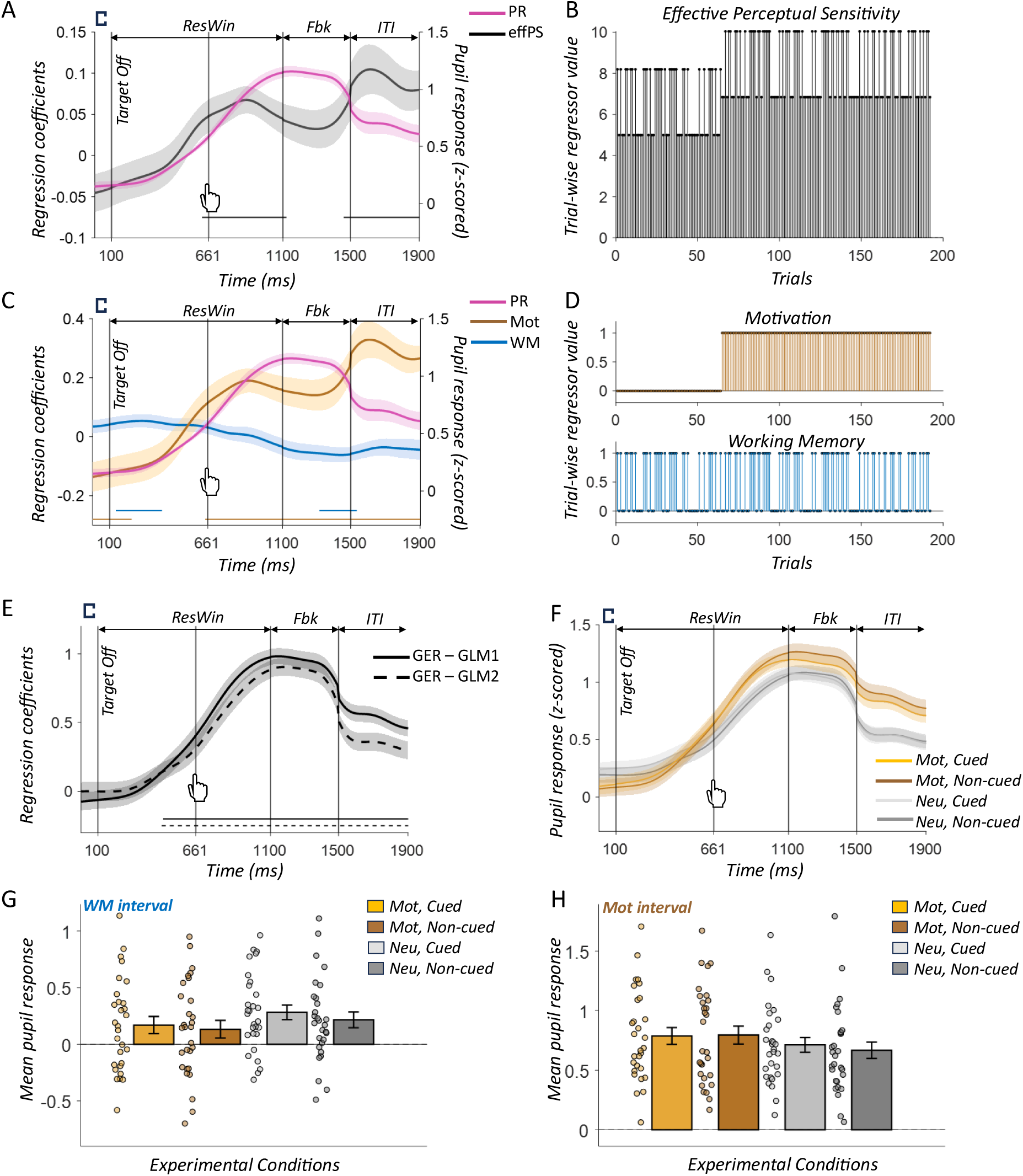
upil dynamics track effective perceptual sensitivity and reveal temporally dissociable influences of working memory and motivation in the perception window. **(A)** Mean pupil response (PR; magenta, right axis) and time-resolved regression coefficients across participants for effective perceptual sensitivity (effPS; black, left axis) obtained from GLM-1. **(B)** Representative trial-wise effective perceptual sensitivity (effPS) values from a single participant. Trial-wise effPS estimates were derived from the winning computational model and used as the parametric regressor in GLM-1. **(C)** Mean pupil response (PR; magenta, right axis) and time-resolved regression coefficients across participants for the working-memory (WM; blue, left axis) and motivational-state (Mot; brown, left axis) regressors obtained from GLM-2. **(D)** Representative trial-wise working memory (WM) and motivational state (Mot) regressors from a single participant. The WM regressor was coded as 1 for cued trials and 0 for non-cued trials, whereas the motivation regressor was coded as 1 for motivational (gain/loss) trials and 0 for neutral trials. **(E)** Time-resolved regression coefficients across participants for the general event-related (GER) regressor obtained from GLM-1 (solid black) and GLM-2 (dashed black). The GER regressor captured pupil responses associated with task events common across trials, including target presentation, response execution, and feedback presentation. In panels **A**, **C**, and **E**, horizontal significance bars indicate time intervals during which effPS regression coefficients differed significantly from zero following cluster-based permutation correction (p < 0.05, corrected). **(F)** Mean pupil responses across participants for the four experimental conditions of the reduced 2 × 2 design: {*Motivational, Cued*} (light brown), {*Motivational, on cued*} (dark brown), { *eutral, Cued*} (light grey), and { *eutral, on cued*} (dark grey). In panels **A**, **C**, **E**, and **F**, shaded regions indicate ± s.e.m. across participants and vertical dashed lines indicate the following events: target offset (100 ms), mean response time (RT = 661 ms), feedback onset (Fbk; 1100 ms), and the onset of the inter-trial interval (ITI; 1500 ms). The response window (ResWin; 100–1100 ms) spans the period between target offset and feedback onset. **(G)** Condition-wise mean pupil responses across participants averaged over time points of the response window, showing significant WM-related effect in GLM-2 at group level (termed as the WM interval), shown separately for the four experimental conditions. **(H)** Condition-wise mean pupil responses across participants averaged over time points of the response window, showing significant motivation-related effect in GLM-2 at group level (termed as the Mot interval). In panels G and H, bars represent group means, error bars indicate ± s.e.m., and markers represent individual participant data points *= 9* .

The GER regressor exhibited significant effects on pupil responses across most of the perception window, beginning after target offset and extending through response execution and feedback processing (475–1900 ms), consistent with task-evoked pupil responses associated with common trial events (see horizontal significance bar in **Fig. 4E**; all p < 0.05 corrected for multiple comparisons across time using cluster-based permutation approach, see *Methods*). After accounting for general event-related pupil responses, effPS significantly modulated pupil dynamics during two distinct intervals within the perception window (all p < 0.05, corrected for multiple comparisons; **Fig. 4A**). The first significant interval emerged around response execution, i.e., during the late stages of response formation and extended into the post-response period (625–1116 ms). Because this interval spanned response execution and preceded feedback onset, the observed modulation suggests that pupil dynamics tracked trial-by-trial fluctuations in processes contributing to perceptual decision formation. The persistence of this effect into the post-response period further suggests that pupil dynamics continued to reflect processing related to the completed perceptual judgment while participants awaited feedback.

A second significant interval was observed near the end of the feedback period (1450–1900 ms; **Fig. 4A**). Because feedback is evaluated relative to the perceptual judgment that immediately preceded it, trial-by-trial variations in effPS continued to modulate pupil dynamics during outcome processing. Given the relatively slow temporal dynamics of pupil responses, this late effect likely reflects the continued influence of perceptual sensitivity during feedback evaluation, indicating that its impact extended beyond response formation and into the processing of behavioural outcomes.

Together, these findings demonstrate that, beyond the robust pupil responses associated with general task events, pupil dynamics reliably tracked latent trial-by-trial fluctuations in effective perceptual sensitivity during response formation and feedback evaluation periods. This pattern indicates that the influence of perceptual sensitivity extended beyond perceptual judgment itself and persisted during the processing of its outcome.

### Temporal dissociation of working-memory and motivational influences on pupil responses

To dissociate the individual contributions of working memory and motivation to pupil dynamics, we performed a second time-resolved GLM (GLM-2) including separate regressors for experimental variables working-memory (*W_t_*) and motivation (*M_t_*) in addition to the GER regressor (see *Methods*; **Fig. 4C-E**). **Figure 4D** shows representative trial-wise values of regressors motivation and WM for a single participant. As in GLM-1, the GER regressor exhibited significant effects on pupil responses across most of the perception window in GLM-2 as well (466–1900 ms, all p < 0.05 corrected for multiple comparisons; **Fig 4E**).

Beyond these general task-evoked effects, the second GLM revealed temporally dissociable effects of working memory and motivation across both the response and feedback periods (all p < 0.05, corrected for multiple comparisons; see horizontal significance bars in **Fig. 4C**). During response window, WM-related effects emerged shortly after target offset and persisted into the early response window (133–400 ms), suggesting that maintained WM representations influenced perceptual decision formation prior to response execution. Notably, WM-related modulation re-emerged during the later stages of feedback presentation (1308–1525 ms), indicating that the influence of the maintained memory representation persisted beyond decision formation and continued to shape the evaluation of behavioural outcomes. The early motivation-related effect emerged during target presentation and persisted beyond target offset (0–225 ms), suggesting that motivational state influenced both the initial encoding of sensory information and the subsequent processing of the perceptual representation. This temporal profile is consistent with motivation enhancing task engagement and the utilization of sensory evidence during the early stages of perceptual decision formation. A second motivation-related effect emerged around the time of response execution and persisted continuously throughout the post-response and feedback periods until the end of the trial (650–1900 ms), indicating a sustained influence of motivational state on post-decisional processing, anticipation of behavioural outcomes, and feedback evaluation.

Further, to directly quantify the condition-wise differences underlying the WM- and motivation-related pupil effects identified by the second GLM, we performed separate 2 × 2 repeated-measures ANOVAs on mean pupil responses extracted from the WM and motivation intervals of the response window (see *Methods*; **Fig. 4F-H**). Analyses were restricted to the response window because it most directly captured the influence of working memory and motivation on perceptual decision formation and behavioural performance. Within the WM interval, the ANOVA revealed a significant main effect of WM state (*F*(1,28) 5.33, *p* 0.03), with larger pupil responses observed for cued relative to non- cued trials. Neither the main effect of Motivation (*F*(1,28) 3.54, *p* 0.08) nor the WM × Motivation interaction (*F*(1,28) 0.35, *p* 0.56) reached significance, indicating that the observed pupil modulation within WM interval was selectively driven by WM cueing. In contrast, within the motivation interval, the ANOVA revealed a significant main effect of Motivation (*F*(1,28) 6.84, *p* 0.01), with larger pupil responses observed during motivational relative to neutral trials. Neither the main effect of WM state (*F*(1,28) 0.65, *p* 0.43) nor the interaction (*F*(1,28) 1.16, *p* 0.29) was significant, indicating that pupil modulation within Mot interval was selectively driven by motivational state. These findings complement the time-resolved GLM results by demonstrating that the early WM-related pupil effect primarily reflected differences between cued and non-cued trials, whereas the later motivation-related effect primarily reflected differences between motivational and neutral trials. Importantly, the absence of significant effects of motivation within the WM interval, and of WM within the motivation interval, confirms that the two temporal windows selectively captured WM- and motivation-related influences on pupil dynamics, respectively.

Overall, the behavioural, computational, and pupillometric findings converge to demonstrate temporally dissociable influences of working memory and motivation on perceptual performance. Working memory cueing improved accuracy, reduced response times, and elicited pupil modulation shortly after target onset in the early response window, suggesting that maintained WM representations rapidly biased the processing of target stimulus at the remembered spatial location, thereby facilitating perceptual decision formation prior to response execution. In contrast, motivation improved performance while exerting a more sustained influence on pupil dynamics extending from early stimulus processing through response execution to outcome evaluation, enhanced both the utilization of sensory evidence during perceptual decision making and the processing of behaviourally relevant outcomes during later stages of the trial. Together, these findings indicate that working memory primarily facilitated perceptual decision formation, whereas motivation exerted a broader influence spanning both decisional and post-decisional processes.

## Discussion

The present study investigated how motivational state and visuospatial working memory jointly shape perceptual sensitivity by integrating behavioural, computational, and physiological approaches. Although previous studies have independently demonstrated beneficial effects of WM and motivation on perception, the mechanisms through which these internal states interact during perceptual processing remained unclear. Two competing possibilities were considered: motivation may amplify WM-driven facilitation to modulate perceptual sensitivity, or motivation and WM may independently contribute to perceptual sensitivity. Across behavioural, computational, and pupillometric analyses, our findings consistently supported the latter hypothesis. Behaviourally, both motivation and WM independently enhanced perceptual discrimination, producing the greatest improvement when both factors were simultaneously present. Computational modelling further demonstrated that these enhancements were best explained by independent additive contributions of motivation and WM to effective perceptual sensitivity rather than motivational amplification of WM-driven facilitation. Finally, pupil dynamics tracked model-derived fluctuations in effective perceptual sensitivity while revealing temporally dissociable signatures of WM and motivation during perceptual decision making. Together, these findings provide converging evidence that multiple top-down cognitive systems independently enhance perceptual sensitivity through complementary computational mechanisms.

Our behavioural findings extend previous demonstrations that WM facilitates visual perception. Numerous studies have shown that information actively maintained in WM biases attention toward memory-matching stimuli and improves perceptual discrimination ^3,4,9,11^. However, much of this work employed feature-based memory representations or visual search paradigms containing multiple competing stimuli, making it difficult to distinguish genuine enhancement of sensory representations from improved distractor filtering ^7,10^. By using a single near-threshold Landolt stimulus without competing distractors, the present study minimized stimulus competition and therefore provided stronger evidence that active visuospatial WM directly enhanced perceptual sensitivity. Furthermore, the comparable behavioural performance observed for uncued and mismatched trials suggests that simple residual activation arising from previous cue presentation was insufficient to facilitate perception ^11^. Instead, the superior performance observed in cued trials is more consistent with active maintenance of task-relevant spatial information rather than passive priming, supporting recent proposals that WM selectively biases sensory processing through actively maintained representations rather than residual stimulus activation ^5,11^.

The behavioural results also demonstrate that motivational incentives independently improved perceptual performance. Both gain and loss contexts enhanced discrimination accuracy and efficiency relative to neutral trials, consistent with previous reports that motivational significance enhances perceptual processing and increases behavioural sensitivity^16,18,19^. Importantly, gain and loss produced statistically comparable improvements, suggesting that motivational salience, rather than reward valence per se, was sufficient to facilitate perceptual performance. More critically, behavioural measures revealed no evidence that motivation selectively enhanced WM-driven facilitation. Instead, motivation and WM contributed independently to behavioural improvements, with the combined motivational and cued condition producing the highest performance. Although reaction times showed a significant interaction, this pattern likely reflects reduced opportunity for further response-speed improvements when perceptual decisions had already been facilitated by either WM or motivational state. Consistent with this interpretation, the primary behavioural measures of efficiency score and accuracy both supported additive rather than synergistic influences of motivation and WM.

The computational modelling provides a mechanistic explanation for these behavioural observations. Rather than relying solely on descriptive behavioural measures, we estimated trial-by-trial fluctuations in effective perceptual sensitivity, a latent variable representing the computational gain of perceptual processing. Critically, the nested model comparison directly evaluated the two competing hypotheses proposed in the Introduction. The winning model incorporated baseline perceptual sensitivity, WM-driven facilitation, and direct motivational modulation of perception while excluding motivational modulation of WM-driven facilitation. Thus, the data provided positive evidence that motivation enhances perceptual processing through a global increase in perceptual sensitivity rather than by selectively strengthening WM-related facilitation. This distinction is theoretically important because a nonsignificant behavioural interaction alone cannot discriminate between alternative computational architectures ^23,34,35^. In contrast, Bayesian model comparison explicitly demonstrated that independent additive pathways better explained behavioural performance than models proposing hierarchical modulation of WM by motivation. The high protected exceedance probability, successful leave-one-participant-out prediction, and strong correlations between effective perceptual sensitivity and behavioural measures further indicate that the model captured a meaningful latent mechanism underlying perceptual decision making rather than simply fitting observed behaviour.

These findings contribute to growing evidence that multiple top-down cognitive processes converge on sensory processing through distinct computational pathways. Working memory has been proposed to selectively enhance representations of behaviourally relevant information by increasing the priority of maintained sensory templates ^1,6^, whereas motivational state has been suggested to globally increase behavioural readiness and sensory gain through value-dependent mechanisms ^18,36^. Our computational results support this conceptual distinction. Whereas WM selectively enhanced processing of stimuli appearing at the maintained spatial location, motivational state produced a broader enhancement of perceptual sensitivity that operated independently of WM state. Thus, rather than interacting within a single hierarchical mechanism, WM and motivation appear to represent complementary top-down systems whose effects converge at the level of perceptual sensitivity.

The pupillometry results provide an important physiological validation of this computational framework. Pupil diameter has been widely associated with cognitive effort, arousal, and locus coeruleus-noradrenergic activity ^24,25,37,38^. Our first GLM demonstrated that pupil responses tracked trial-by-trial fluctuations in model-derived effective perceptual sensitivity after accounting for general task-evoked responses. This finding suggests that pupil dynamics reflected the latent computational variable underlying perceptual performance rather than merely sensory stimulation or motor execution. The persistence of this relationship during both response formation and feedback processing further indicates that effective perceptual sensitivity continued to influence post-decisional evaluation, consistent with the relatively slow temporal dynamics of the pupil response.

By dissociating WM and motivation within a second GLM, we further demonstrated distinct temporal signatures of these top-down processes. WM-related pupil modulation emerged shortly after target presentation and during the early response window, suggesting rapid prioritization of maintained spatial representations during perceptual decision formation. In contrast, motivational influences appeared during initial stimulus processing and persisted throughout response execution and feedback evaluation, indicating a broader role in sustaining task engagement, utilization of sensory evidence, and processing of behavioural outcomes. The accompanying ANOVAs confirmed that these temporal intervals selectively reflected WM and motivational influences, respectively, with little evidence for cross-factor modulation. These results complement the computational findings by demonstrating that, although WM and motivation converge on the common outcome of enhanced perceptual sensitivity, they remain temporally dissociable physiological processes.

Several limitations should be acknowledged. First, the study examined healthy young adults performing a visuospatial discrimination task with monetary incentives, and therefore generalization to other age groups, sensory modalities, or motivational contexts requires further investigation. Second, although pupillometry provides a valuable index of cognitive and arousal-related processes, it does not directly identify the underlying neural circuits. Future studies combining computational modelling with electrophysiological or neuroimaging approaches could determine how frontoparietal control systems, visual cortex, and neuromodulatory networks jointly implement the independent influences of WM and motivation identified here. Finally, extending the computational framework to more complex naturalistic environments may further clarify how multiple cognitive states dynamically regulate perception during real-world behaviour.

In conclusion, the present study demonstrates that visuospatial working memory and motivational state independently enhance perceptual sensitivity through complementary computational mechanisms. Behavioural performance, computational modelling, and pupillometry converged on the conclusion that motivation does not amplify WM-driven facilitation but instead contributes an independent increase in perceptual sensitivity while exhibiting a distinct temporal profile during perceptual decision making. By integrating behavioural, computational, and physiological evidence within a unified framework, this work advances our understanding of how multiple top-down cognitive systems cooperate to optimize perception under goal-directed behaviour and provides a mechanistic foundation for future investigations of perceptual dysfunction in conditions involving impairments of cognition and motivation.

## Methods

### Participants

A sample size of 32 participants was based on statistical power analysis conducted using G*Power tool ^39^ for a within-subjects 3 × 3 repeated-measures design, with factors motivational state (loss, gain, neutral) and WM location during target presentation (cued, uncued, mismatched). The participants were enrolled through an online recruitment system. All participants reported normal or corrected-to-normal vision, no history of neurological or psychiatric disorders, and performed the experiment using their dominant hand. All participants were naïve to the hypotheses of the study. Participants received a fixed monetary compensation for participation (₹120 per hour) along with a performance-dependent bonus tied to accuracy in the main task. All participants provided written informed consent prior to the experiment and participated in a practice session. The study was approved by the Institute Ethics Committee, Indian Institute of Technology Jodhpur under reference number IEC/IITJ/ 2023-24/9.24.

For behavioural analyses, data from one participant was excluded due to poor perceptual discrimination performance in the neutral baseline condition (38.5% accuracy), falling below the predefined acceptable limit (70% target threshold ± 20% tolerance). The final behavioural sample therefore comprised 31 participants (age: 23.07 ± 3.50 years old; 15 females). For eye-tracking (pupillometry) analyses, two more participants were excluded due to data loss (greater than 20% of trials lost due to excessive blinking), in addition to the participant excluded from behavioural analyses. The final eye-tracking sample thus consisted of 29 participants (age: 22.90 ± 3.50 years old; 14 females).

### Experimental Design

#### Stimuli

Visual stimuli corresponding to distinct task components, included motivational, visuospatial working memory, and perceptual target stimuli (**Fig. 1A, B**). Stimuli were presented against a uniform grey background on a calibrated monitor (resolution 1,080 × 1,920 pixels, refresh rate = 120 Hz), positioned at a 60 cm viewing distance. Monetary associations (motivational state) in the main task were conveyed via colour of the central fixation square (0.5° visual angle) presented throughout each trial, with grey indicating neutral (no monetary outcome), green indicating reward (gain), and red indicating penalty (loss) conditions. In gain trials, participants earned ₹10 for each correct response and received no money for incorrect responses. In loss trials, participants incurred a penalty of −₹10 for each incorrect response, with no money for correct responses. At the end of the session, participants received their total earnings based on task performance (up to a maximum of ₹640), in addition to a fixed participation fee of ₹120 per hour. Visuospatial working memory stimuli consisted of coloured circular cues (0.6° diameter) presented at 16 equidistant peripheral locations arranged along an imaginary circle centred on fixation at 6° eccentricity (radial distance from central fixation square), spanning the full peripheral visual field ^40^. Cue colours differed across trial types, with blue used for the main trials and orange for memory probe trials. The perceptual target employed was a Landolt stimulus (0.5° visual angle), a standardized shape with a small gap (0.15° visual angle) appearing as a broken shape, based on a previous work ^11^. Participants were required to identify the position of the gap (left or right), allowing precise assessment of visual discrimination performance. The Landolt stimulus was always presented on top of a coloured circular cue at one of 16 peripheral locations arranged around fixation. Thus, the target and the cue were spatially aligned, appearing at the same location on each trial (**Fig. 1A, B**).

### Perceptual threshold task

Prior to the main task, each participant performed a perceptual threshold task, to estimate a participant-specific target presentation duration at an accuracy of 70% (**Fig. S1**). Participants maintained central fixation throughout the threshold task. On each trial, they performed a visual discrimination task, reporting the gap side (left or right) of the Landolt stimulus presented on top of a blue circular cue at one of the 16 peripheral locations. Each trial began with a jittered fixation period (2000 ± 460 ms), followed by a brief target presentation (see next paragraph for duration). Participants responded within 1000 *ms* of target offset, after which feedback was provided (400 ms; correct/incorrect). The threshold task comprised of 192 trials in total. The sixteen spatial locations and the two gap sides (left, right) were equally represented and presented in pseudo-random order. Gap direction was balanced across spatial locations and hemispheres. Short breaks (≈60 s) were provided after every 16 trials to minimize fatigue.

Target presentation duration in each trial was adaptively manipulated using a double random interleaved staircase procedure ^30^ implemented via two overlapping duration ranges: staircase A (100, 120, 140 *ms*) and staircase B (140, 160, 180 *ms*). The double interleaved staircase method estimates perceptual threshold by running two independent adaptive staircases in parallel, with trials randomly drawn from either staircase. Each staircase varies stimulus intensity (here, target duration) based on performance. Random interleaving prevents participants from predicting trial difficulty, thereby minimizing expectancy effects and response strategies. The durations in staircases A and B were selected to cover the range around near-threshold discriminability (70%) for brief peripheral presentations at ∼6° eccentricity, where visual discrimination is constrained by reduced acuity ^40^. In particular, the duration range (100-180 ms) was selected based on pilot testing to span performance from near-chance to saturation levels (∼50-100%) for the present peripheral stimulus configuration. The two staircases were designed to overlap at 140 ms. This overlap allowed both staircases to converge toward a common performance level while reducing sensitivity to starting values and improving the stability of threshold estimation.

Within each staircase, a one-up/three-down rule ^41^ was applied, where a single incorrect response increased stimulus duration (easier), whereas three consecutive correct responses decreased duration (harder). This asymmetric rule allowed convergence at approximately 70% correct performance, as decreases in duration require sustained accuracy, whereas increases occur after a single error. Consequently, not all stimulus durations necessarily occurred for every participant. For example, if three consecutive correct responses were not achieved at a given level (e.g., 120 ms), the staircase did not progress to shorter durations (e.g., 100 ms). On the other hand, consistently correct responses at a given duration (e.g., 160 *ms*) prevented progression to longer durations (e.g., 180 ms). Thus, the effective range of presented durations was participant-specific.

Performance across durations was fit for each participant using a sigmoidal psychometric function characterized by bias (*μ*) and sensitivity (*σ*) parameters (**Fig. S1**). The bias parameter (*μ*) reflects the point of subjective equality, whereas the sensitivity parameter (*σ*) indexes the steepness of the psychometric curve, i.e., sensitivity to changes in stimulus duration. The fitted function was then used to estimate the target presentation duration corresponding to 70% discrimination accuracy for each participant. By combining two interleaved staircases with adaptive updating and psychometric fitting, the procedure yielded a robust and unbiased estimate of participant-specific perceptual threshold duration.

### Main task

The main task combined motivational and visuospatial working memory state manipulations to examine their independent and joint effects on visual perception (**Fig. 1A, B**). To ensure compliance with the WM requirement, memory probe trials were interleaved throughout the task. Each participant completed 240 trials, comprising 192 main task trials and 48 memory probe trials. Short breaks (≈60 s) were provided after every 40 trials to minimize fatigue.

The main task consisted of trials varying across motivational state (neutral, gain, loss) and WM condition (cued, non-cued). The task began with 64 neutral trials, consisting of 32 cued and 32 non-cued trials (equally divided into uncued and mismatched conditions) presented in a pseudo-random order. This was followed by 64 loss and 64 gain trials, also presented in pseudo-random order. Within each motivational condition (loss and gain), trials were evenly split between cued and non-cued trial types. A cued trial was defined as one in which the perceptual target appeared at the spatial location actively maintained in working memory, whereas a non-cued trial was one in which the target appeared at a different location (see next paragraph for definitions of uncued and mismatched trials). Neutral trials preceded motivational trials to establish an unbiased baseline and minimize carryover effects of motivational state due to gain or loss. Additionally, 16 memory probe trials were interleaved within each main task condition (neutral, gain, loss), yielding a total of 48 memory trials.

Participants maintained central fixation throughout the experiment (**Fig. 1A, B**). Trials were separated by a jittered inter-trial interval (ITI; 2000 ± 460 ms). Each trial began with the sequential presentation of two blue circular WM cues (500 ms each) separated by a 1000 *ms* interval. The cue positions were distinct and drawn from 16 equidistant peripheral locations. A retro-cue (1 or 2; 500 ms) then indicated which location was to be maintained in WM. After a jittered delay (4000 ± 650 ms), a Landolt target briefly appeared according to participant-specific threshold duration (range: 100-180 ms) at either the cued, uncued, or mismatched spatial location. The cued location corresponded to the retro-cued position, the uncued location to a presented position but unselected for maintenance, and the mismatched location to a position not presented during sequential cue presentation period. Participants reported the gap side (left/right) within 1000 ms of target offset, followed by feedback (correct/incorrect; 400 ms).

Memory probe trials followed the same trial sequence as main trials but omitted the perceptual target. Instead, participants judged whether a test location matched the retro-cued location. The test location could be cued, uncued, or mismatched, and these trials were distinguished by orange cues (**Fig. 1A, B**).

The sixteen spatial locations, two retro-cue numbers (1, 2), and two target gap sides (left, right) were equally represented across all conditions of the 3 × 3 design (cued, uncued, mismatched × neutral, gain, loss) as well as in the memory probe trials. Each spatial location appeared equally often as Cue 1, Cue 2, and as a target location, and was retro-cued an equal number of times. Gap direction (left/right) was balanced across spatial locations and hemispheres.

### Computational model of perceptual decision behaviour

To provide a **mechanistic account of perceptual performance**, behavioural responses were modelled using Bayesian generalized linear framework grounded within psychometric decision theory, here referred to as a generalized linear psychometric model ^23^. This approach was inspired by Bayesian generalized linear mixed modelling ^42^ and ideal observer analysis in signal detection theory ^43,44^. For broader methodological overviews, refer elsewhere ^23,35,45^. Responses were modelled as binary outcomes (correct vs. incorrect) under a logistic decision function. For participant *i*, the probability of a correct response on trial *t* was determined by a latent decision variable, corresponding to the slope of the decision function:

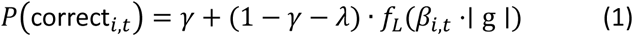

Where, *γ* and *λ* are non-sensory lower and upper asymptotic lapse rates, respectively, logistic function *f_L_*(*x*) ;1/(1 *e*^−*x*^), *g* denotes the absolute gap width representing stimulus strength (fixed here at *g* 0.15^∘^) and 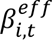 denotes the effective perceptual sensitivity corresponding to the slope of the psychometric decision function. The lower and upper asymptotic error rates were tied together to reduce model complexity and enforce unbiased lapse behaviour ^38^.

For participant *i*, effective sensitivity on trial *t* was modelled as a linear weighted combination of sensory evidence, working memory, and motivational state ^23^:

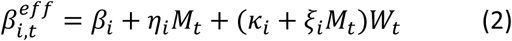

where, coefficients *β_i_*, *κ_i_*, *η_i_*, and *ξ_i_* denote participant-specific strengths of baseline perceptual sensitivity, WM-driven facilitation, direct motivational modulation, and indirect motivational modulation of WM-driven facilitation, respectively. In contrast, *M_t_* and *W_t_* denote trial-wise experimental conditions shared across participants. *M_t_* was coded as 1 for motivational (gain/loss) trials and 0 for neutral trials, whereas *W_t_* was coded as 1 for cued trials and 0 for uncued and mismatched trials. The binary coding of experimental conditions was based on the reduced 2 x 2 condition space and effects observed in the behavioural ANOVA analyses.

This formulation was based on evidence suggesting that motivational states exert a relatively global enhancement on perceptual and cognitive processing ^36^, whereas WM selectively facilitates processing of task-relevant stimuli ^12,14^. Accordingly, the term *η_i_M_t_* models the direct global influence of motivation on perceptual sensitivity independent of WM state, while the term (*κ_i_ ξ_i_M_t_*)*W_t_* captures both the direct WM-driven facilitation of perception (*κ_i_W_t_*) and the indirect motivational amplification of WM-driven facilitation (*ξ_i_M_t_W_t_*). Together, these components determine trial-by-trial fluctuations in effective perceptual sensitivity underlying perceptual discrimination performance (**Fig. 1C**).

### Model comparison

To identify how distinct components contribute to enhanced discrimination accuracy, a family of nested models was constructed from the full model (Eq. 2) by systematically including or excluding (i) direct motivational influences on perceptual sensitivity (*Mot* → *Per*), (ii) WM–driven facilitation effects (*WM* → *Per*), and (iii) motivational modulation of WM-driven facilitation (*Mot* → *WM*). For a similar approach to constructing and testing nested computational models, see Park et al. ^34^. This procedure yielded a total of eight candidate models, including the full model and a null model capturing only baseline perceptual sensitivity without contributions from motivation or working memory (Fig. 3A).

The candidate models were then compared using random-effects Bayesian model comparison implemented using the Variational Bayesian Analysis (VBA) toolbox in MATLAB ^33^. First, at the subject level, participant-specific parameters (*β_i_*, *κ_i_*, *η_i_*, *ξ_i_*) were estimated independently using maximum a posteriori (MAP) estimation with weakly informative Gaussian priors to minimize undue prior influence on posterior estimates ^34^. Priors were defined in unconstrained parameter space with mean (*μ*=0) and large variance (*σ*=5), based on previous work ^34^. Model parameters (*β_i_*, *κ_i_*, *η_i_*, *ξ_i_*) were constrained to be positive values within a range (0−5). Upper parameter bounds were selected to remain within biologically plausible ranges of increases in psychometric slope and perceptual sensitivity produced by top-down modulation in visual psychophysics ^46^.

For each participant, model fit was quantified using the negative log posterior at the MAP solution, corresponding to a model evidence estimate incorporating both likelihood and prior terms ^47^. These subject-wise model evidence estimates were subsequently entered into a group-level random-effects (RFX) Bayesian model comparison ^32^. This hierarchical RFX formulation allows for inter-individual variability, such that different participants may be better explained by different models rather than enforcing a single best model across all participants. Group-level inference was then performed over the posterior distribution of model evidences, from which expected model frequencies and protected exceedance probabilities were derived. Expected model frequency represents the estimated proportion of participants in the population for whom a given model is most likely to explain behavioural data under the random-effects Bayesian framework. The winning model was identified as the model with the highest protected exceedance probability. Protected exceedance probability is derived from the posterior distribution of expected model frequencies and quantifies the probability that a given model is more frequent than all competing models in the population beyond chance-level differences ^32,48^.

### Pupil data acquisition and preprocessing

Pupil diameter from the left eye was recorded using a Tobii Pro Fusion eye tracker (Tobii AB, Stockholm, Sweden) at a sampling rate of 120 Hz. A 9-point calibration procedure was employed at the start of the main task and after each break during the task. A chin rest was used throughout the experiment to reduce head movements and maintain stable eye-tracking quality. Saccades and blinks were detected using custom routines for preprocessing eye-tracking data implemented in MATLAB (version 24.1; The MathWorks, Natick, MA, USA). To maintain central fixation throughout the experiment, gaze position was continuously monitored online. Whenever participants deviated by more than 1° of visual angle from the fixation point, the central fixation square increased in size, providing immediate visual feedback to guide gaze back to fixation. Periods of signal loss due to blinks were reconstructed using linear interpolation with a half-window of 50 ms before and after each blink. The interpolated signal was subsequently low-pass filtered at 4 Hz using a third-order Butterworth filter. Participants for whom more than 20% of total trials were lost due to signal loss or no response, were removed from further analysis (percentage removed trials across 29 participants, mean ± SD = 3.72% ± 5.78%)

Pupil diameter reflects both slower tonic changes associated with arousal and motivation, as well as transient phasic responses linked to task-related cognitive processes ^25,37,49^. Consequently, global z-scoring across all trials may be influenced by condition-dependent tonic shifts, potentially reducing sensitivity to transient WM-related effects. To preserve sensitivity to WM effects within each motivational state, pupil data were normalized separately for neutral and motivational conditions. Specifically, cued and non-cued trials were z-scored together within the neutral condition and separately within the motivational (gain and loss) conditions using the corresponding participant-specific mean and standard deviation. This approach reduced the influence of broad motivational differences in baseline pupil diameter while preserving WM-related fluctuations within each motivational state ^24,25^.

### Pupil regression analysis

To examine whether pupil dynamics tracked computational and experimental factors underlying perceptual performance, we employed two complementary time-resolved general linear model (GLM) analyses during the perception window (target onset to post-feedback; **Fig. 1A**) (Palaskar & Dang, 2024). The first GLM examined whether pupil responses tracked trial-by-trial fluctuations in effective perceptual sensitivity (effPS) estimated from the winning computational model. Because effPS represents an integrated measure combining the influences of working-memory and motivation on perceptual sensitivity, a second GLM was designed to temporally dissociate the independent contributions of working-memory and motivational state to pupil dynamics by modelling their trial-wise experimental conditions separately. Together, these analyses allowed us to determine whether pupil responses reflected the integrated computational variable underlying perceptual performance (effPS) and to identify the distinct temporal signatures of working memory and motivation contributing to that variable (**Fig. 4**).

For the first GLM, trial-wise estimates of effPS were derived from the winning model identified through the nested model comparison procedure described above and computed using participant-specific parameters (*β_i_*, *κ_i_*, *η_i_*) and trial-wise experimental conditions (*W_t_*, *M_t_*). These estimates reflected trial-by-trial variations in perceptual sensitivity arising from the combination of baseline perceptual sensitivity (*β_i_*), WM-driven facilitation (*κ_i_W_t_*), and direct motivational modulation (*η_i_M_t_*). For each participant, GLM analysis was performed on preprocessed, z-scored pupil time series from the main task. At each time point within a trial, pupil responses were modelled using two regressors: (i) a general event-related (GER) regressor capturing pupil responses associated with task events common to all trials, including target presentation, motor response, and feedback presentation, and (ii) a parametric effPS regressor indexing trial-by-trial fluctuations in model-derived perceptual sensitivity (z-scored within participants; see representative trial-wise effPS values for a single participant in **Fig. 4B**). The GER regressor was coded as 1 during the target, response, and feedback epochs and 0 otherwise, thereby accounting for pupil responses associated with task events common across trials. The effPS regressor captured latent variations in effective perceptual sensitivity estimated from the winning computational model. Both regressors were entered simultaneously into the GLM to dissociate sensitivity-related pupil modulation from general event-related pupil responses.

For the second GLM, pupil responses were modelled using three regressors: (i) a general event-related (GER) regressor as for the first GLM, (ii) a working-memory regressor (*W_t_*; coded 1 for cued and 0 for non-cued trials), and (iii) a motivational-state regressor (*M_t_*; coded 1 for motivational and 0 for neutral trials). All regressors were entered simultaneously into the model. This analysis enabled temporal dissociation of WM-related and motivational contributions to pupil dynamics that were otherwise combined within the integrated effPS measure (see representative trial-wise *W_t_* and *M_t_* regressors for a single participant in **Fig. 4D**).

Regression coefficients (beta weights) were estimated separately at each time point relative to target onset, yielding time-resolved measures of the contribution of each regressor to pupil dynamics. For group-level inference, a random-effects analysis was performed using a summary-statistic approach, whereby participant-specific regression coefficients were entered into one-sample t-tests against zero at each time point (Penny and Holmes, 2003). Multiple-comparison correction across time was performed using a non-parametric cluster-based permutation approach. Contiguous time points exceeding an initial uncorrected threshold (p < 0.05) were identified as candidate clusters, and each cluster was assigned a statistic equal to the sum of its constituent t-values. The significance of the observed clusters was determined by comparison with a null distribution generated from 1000 permutations, considered significant at p < 0.05 (cluster-corrected); shown as horizontal significance bars in **Fig. 4A, C, E**).

### ANOVA on mean pupil responses

To complement the second time-resolved GLM analysis, we conducted an additional ANOVA on mean pupil responses (**Fig. 4F-H**). Whereas the GLM analysis were designed to identify the temporal relationship between pupil dynamics and experimental variables, it did not directly quantify condition-wise differences in pupil dynamics across the 2 × 2 experimental design: {Motivation, Cued}, {Motivation, Non-cued}, {Neutral, Cued}, and {Neutral, Non-cued}. Therefore, an additional analysis was performed to examine how pupil responses differed across conditions as a function of working-memory state and motivational state. For each participant, pupil responses were averaged separately within two intervals identified using the second GLM analysis: the WM interval (defined by the significant *W_t_* effect) and the motivation interval (defined by the significant *M_t_* effect) within the response window (see *upil regression analysis*). Because these intervals most directly captured the influence of working memory and motivation on perceptual decision formation and performance, subsequent analyses were restricted to these response window intervals. This yielded mean pupil responses for each of the four experimental conditions. The resulting values were entered into separate 2 × 2 repeated-measures ANOVAs with within-subject factors Motivation (motivational, neutral) and WM (cued, non-cued) to determine whether pupil modulation within each interval of the response window was selectively associated with its corresponding factor or additionally influenced by the other factor and their interaction. Significant effects were followed by Bonferroni-corrected post hoc comparisons. This analysis allowed us to characterize the independent and joint effects of working memory and motivation on condition-wise pupil responses.

## Authors’ Contributions

GB and SD conceptualized the project and designed the task. GB conducted the experiments, preprocessed and analyzed the data. GB and SD interpreted the results. GB wrote the first draft of the manuscript. All authors revised the manuscript.

## Data and Code Availability

The data and custom code that support the findings of this study are available from the corresponding author upon reasonable request.

## Supplementary Material

### Perceptual threshold estimation

Participant-specific perceptual thresholds were estimated using a double interleaved staircase procedure combined with psychometric fitting (see *Methods* and **Fig. S1**). **Figure S1A** illustrates the trial sequence of the threshold task, whereas **Fig. S1B** shows a representative psychometric fit from a single participant, demonstrating the increase in discrimination performance with increasing target duration and the extraction of the 70% threshold duration (ThrDur) from the fitted function. Across participants ( = 31), the mean threshold duration corresponding to 70% discrimination accuracy was 142 ± 4 ms (mean ± SEM; **Fig. S1C**). The mean central tendency parameter (*μ*), reflecting the location of the psychometric function along the duration axis, was 120 ± 2 ms, whereas the mean sensitivity parameter (*σ*), indexing sensitivity to changes in stimulus duration, was 26 ± 2 ms. These results indicated reliable estimation of participant-specific perceptual thresholds for use in the main task.

**Figure S1.**
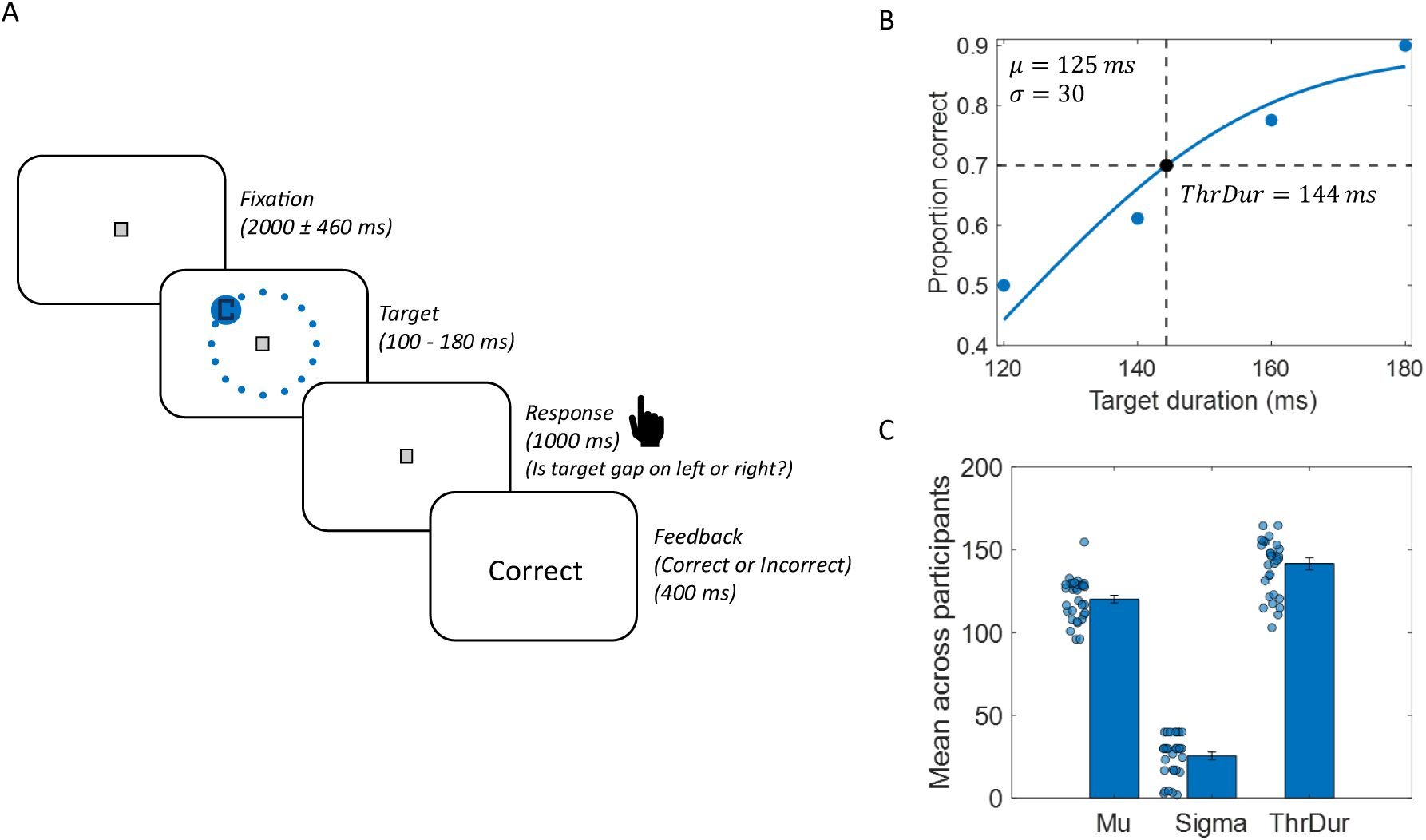
erceptual threshold task and psychometric threshold estimation. **(A)** Trial structure of the perceptual threshold task. Each trial began with a fixation period, followed by brief presentation of the Landolt stimulus superimposed on a blue circular cue at one of 16 peripheral locations. Participants reported the side of the target gap (left/right), followed by performance feedback. Target duration was adaptively varied using a double interleaved staircase procedure. **(B)** Representative psychometric fit (solid blue curve) and observed data points (blue markers) from a single participant, showing proportion correct as a function of target presentation duration. Dashed lines indicate the estimated target duration corresponding to 70% discrimination accuracy (144 ms) extracted from the fitted sigmoidal function, characterized by a central tendency parameter of *μ* 125 *ms* and a sensitivity parameter of *σ* 30 *ms*. **(C)** Group-level mean values (bars; mean ± SEM) and individual participant data points for the psychometric parameters obtained from the threshold task: central tendency parameter (*μ*), sensitivity parameter (*σ*), and the estimated threshold duration (ThrDur; ms) corresponding to 70% discrimination accuracy.

### ANOVA Tables

A 3 x 3 repeated measures ANOVA on efficiency scores for 31 participants with Motivation (loss, gain, neutral) and WM (cued, uncued, mismatched) as within-subject factors:

**Table S1.** Post-hoc comparisons for efficiency score.

| Table S1. Post-hoc comparisons for efficiency score |  |  |  |  |
| --- | --- | --- | --- | --- |
| Factor (Motivation or WM) | Comparison between levels | Difference | Standard error | p-value |
| Loss | Cued - Uncued | 0.19 | 0.06 | $5.58 \times 10^{-3}$ |
| Loss | Cued - Mismatched | 0.25 | 0.06 | $7.87 \times 10^{-4}$ |
| **Loss** | Uncued - Mismatched | 0.06 | 0.05 | 0.51 |
| **Gain** | Cued - Uncued | 0.18 | 0.06 | 0.02 |
| **Gain** | Cued - Mismatched | 0.31 | 0.05 | $1.86 \times 10^{-6}$ |
| **Gain** | Uncued - Mismatched | 0.13 | 0.06 | 0.09 |
| **Neutral** | Cued - Uncued | 0.20 | 0.04 | $1.70 \times 10^{-4}$ |
| **Neutral** | Cued - Mismatched | 0.34 | 0.05 | $9.76 \times 10^{-8}$ |
| **Neutral** | Uncued - Mismatched | 0.14 | 0.05 | 0.02 |
| **Cued** | Loss - Gain | -0.02 | 0.03 | 0.89 |
| **Cued** | Loss - Neutral | 0.19 | 0.05 | $6.16 \times 10^{-4}$ |
| **Cued** | Gain - Neutral | 0.21 | 0.04 | $3.36 \times 10^{-5}$ |
| **Uncued** | Loss - Gain | -0.03 | 0.04 | 0.79 |
| **Uncued** | Loss - Neutral | 0.20 | 0.04 | $9.75 \times 10^{-5}$ |
| **Uncued** | Gain - Neutral | 0.23 | 0.05 | $2.53 \times 10^{-4}$ |
| **Mismatched** | Loss - Gain | 0.05 | 0.05 | 0.55 |
| **Mismatched** | Loss - Neutral | 0.28 | 0.05 | $2.71 \times 10^{-5}$ |
| **Mismatched** | Gain - Neutral | 0.23 | 0.04 | $1.49 \times 10^{-5}$ |

A 2 x 2 repeated measures ANOVA on efficiency score for 31 participants with Motivation (motivation, neutral) and WM (cued, non-cued) as within-subject factors:

**Table S2.** Post-hoc comparisons for efficiency score.

| **Table S2. Post-hoc comparisons for efficiency score** |  |  |  |  |
| --- | --- | --- | --- | --- |
| **Factor (Motivation or WM)** | **Comparison between levels** | **Difference** | **Standard error** | **p-value** |
| Motivation | Cued- Non-cued | 0.23 | 0.04 | $1.05 \times 10^{-5}$ |
| Neutral | Cued- Non-cued | 0.27 | 0.04 | $2.72 \times 10^{-8}$ |
| Cued | Motivation - Neutral | 0.20 | 0.04 | $1.85 \times 10^{-5}$ |
| Non-cued | Motivation - Neutral | 0.24 | 0.03 | $2.84 \times 10^{-9}$ |

A 2 x 2 repeated measures ANOVA on accuracy values for 31 participants with Motivation (motivation, neutral) and WM (cued, non-cued) as within- subject factors:

**Table S3.** Post-hoc comparisons for accuracy.

| **Table S3. Post-hoc comparisons for accuracy** |  |  |  |  |
| --- | --- | --- | --- | --- |
| **Factor (Motivation or WM)** | **Comparison between levels** | **Difference** | **Standard error** | **p-value** |
| Motivation | Cued - Non-cued | 6.20 | 1.95 | $3.45 \times 10^{-3}$ |
| Neutral | Cued - Non-cued | 7.98 | 1.82 | $1.30 \times 10^{-4}$ |
| Cued | Motivation - Neutral | 6.25 | 1.56 | $3.71 \times 10^{-4}$ |
| Non-cued | Motivation - Neutral | 8.03 | 1.51 | $9.56 \times 10^{-6}$ |

A 2 x 2 repeated measures ANOVA on reaction times for 31 participants with Motivation (motivation, neutral) and WM (cued, non-cued) as within-subject factors:

**Table S4.** Post-hoc comparisons for reaction time.

| **Table S4. Post-hoc comparisons for reaction time** |  |  |  |  |
| --- | --- | --- | --- | --- |
| **Factor**<br>**(Motivation or WM)** | **Comparison between**<br>**levels** | **Difference** | **Standard error** | **p-value** |
| Motivation | Cued - Non-cued | -0.04 | $4.85 \times 10^{-3}$ | $6.39 \times 10^{-9}$ |
| Neutral | Cued - Non-cued | -0.06 | 0.01 | $1.97 \times 10^{-7}$ |
| Cued | Motivation - Neutral | -0.02 | 0.01 | $1.75 \times 10^{-3}$ |
| Non-cued | Motivation - Neutral | -0.05 | 0.01 | $3.61 \times 10^{-5}$ |

## References

1. Gilbert, C. D. & Li, W. Top-down influences on visual processing. Nat. Rev. Neurosci. 14, 350–363 (2013).

2. Caras, M. L. & Sanes, D. H. Top-down modulation of sensory cortex gates perceptual learning. Proc. Natl. Acad. Sci. U. S. A. 114, 9972–9977 (2017).

3. Kiyonaga, A. & Egner, T. Working memory as internal attention: Toward an integrative account of internal and external selection processes. Psychon. Bull. Rev. 20, 228–242 (2013).

4. Soto, D., Hodsoll, J., Rotshtein, P. & Humphreys, G. W. Automatic guidance of attention from working memory. Trends Cogn. Sci. 12, 342–348 (2008).

5. Hollingworth, A., Matsukura, M. & Luck, S. J. Visual Working Memory Modulates Rapid Eye Movements to Simple Onset Targets. Psychol. Sci. 24, 790–796 (2013).

6. Schneegans, S., Spencer, J. P., Schöner, G., Hwang, S. & Hollingworth, A. Dynamic interactions between visual working memory and saccade target selection. J. Vis. 14, (2014).

7. Cosman, J. D. & Vecera, S. P. The contents of visual working memory reduce uncertainty during visual search. *Attention, Perception*, Psychophys. 73, 996–1002 (2011).

8. Han, S. W. Working memory contents enhance perception under stimulus-driven competition. Mem. Cogn. 43, 432–440 (2015).

9. Pan, Y., Cheng, Q. P. & Luo, Q. Y. Working memory can enhance unconscious visual perception. Psychon. Bull. Rev. 19, 477–482 (2012).

10. Dosher, B. A. & Lu, Z. L. Noise exclusion in spatial attention. Psychol. Sci. 11, 139–146 (2000).

11. Pan, Y. & Zhang, X. Visual working memory enhances target discrimination accuracy with single-item displays. *Attention, Perception*, Psychophys. 82, 3005–3012 (2020).

12. Soto, D., Wriglesworth, A., Bahrami-Balani, A. & Humphreys, G. W. Working Memory Enhances Visual Perception: Evidence From Signal Detection Analysis. J. Exp. Psychol. Learn. Mem. Cogn. 36, 441–456 (2010).

13. Carrasco, M., Williams, P. E. & Yeshurun, Y. Covert attention increases spatial resolution with or without masks: Support for signal enhancement. J. Vis. 2, 467–479 (2002).

14. Pan, Y., Luo, Q. & Cheng, M. Working memory-driven attention improves spatial resolution: Support for perceptual enhancement. *Attention, Perception*, Psychophys. 78, 1625–1632 (2016).

15. Padmala, S. & Pessoa, L. Motivation versus aversive processing during perception. Emotion 14, 450–454 (2014).

16. Hu, K., Padmala, S. & Pessoa, L. Interactions between reward and threat during visual processing. Neuropsychologia 51, 1763–1772 (2013).

17. Zhang, P. et al. High reward enhances perceptual learning. J. Vis. 18, 1–21 (2018).

18. Serences, J. T. Value-Based Modulations in Human Visual Cortex. Neuron 60, 1169–1181 (2008).

19. Engelmann, J. B., Damaraju, E., Padmala, S. & Pessoa, L. Combined effects of attention and motivation on visual task performance: Transient and sustained motivational effects. Front. Hum. Neurosci. 3, (2009).

20. Cho, Y. T. et al. Reward and loss incentives improve spatial working memory by shaping trial-by-trial posterior frontoparietal signals. Neuroimage 254, (2022).

21. Aul, C. et al. The functional relevance of visuospatial processing speed across the lifespan. Cogn. Res. Princ. Implic. 8, (2023).

22. Trés, E. S. & Brucki, S. M. D. Visuospatial processing: A review from basic to current concepts. Dement. Neuropsychol. 8, 175–181 (2014).

23. Houpt, J. W. & Bittner, J. L. Analyzing thresholds and efficiency with hierarchical Bayesian logistic regression. Vision Res. 148, 49–58 (2018).

24. Kloosterman, N. A. et al. Pupil size tracks perceptual content and surprise. Eur. J. Neurosci. 41, 1068–1078 (2015).

25. Mathôt, S. Pupillometry: Psychology, physiology, and function. J. Cogn. 1, (2018).

26. Purves, D., Monson, B. B., Sundararajan, J. & Wojtach, W. T. How biological vision succeeds in the physical world. Proc. Natl. Acad. Sci. U. S. A. 111, 4750–4755 (2014).

27. Yu, K., Vanpaemel, W., Tuerlinckx, F. & Zaman, J. The probabilistic and dynamic nature of perception in human generalization behavior. iScience 28, (2025).

28. Zhang, M. et al. Exploring the spatial working memory and visual perception in children with autism spectrum disorder and general population with high autism-like traits. PLoS One 15, (2020).

29. Sanna, F., Mahmut, M. K., Loy, F. & Masala, C. Editorial: Sensorial and perceptual dysfunctions as predisposing factors for the onset of depression. Front. Neurosci. 16, (2022).

30. Cornsweet, T. The staircrase-method in psychophysics. Am. J. Psychol. 75, (1962).

31. Bruyer, R. & Brysbaert, M. Combining speed and accuracy in cognitive psychology: Is the inverse efficiency score (IES) a better dependent variable than the mean reaction time (RT) and the percentage of errors (PE)? Psychol. Belg. 51, 5–13 (2011).

32. Stephan, K. E., Penny, W. D., Daunizeau, J., Moran, R. J. & Friston, K. J. Bayesian model selection for group studies. Neuroimage 46, 1004–1017 (2009).

33. Daunizeau, J., Adam, V. & Rigoux, L. VBA: A Probabilistic Treatment of Nonlinear Models for Neurobiological and Behavioural Data. PLoS Comput. Biol. 10, (2014).

34. Park, H., Doh, H., Lee, E., Park, H. & Ahn, W. Y. The neurocognitive role of working memory load when Pavlovian motivational control affects instrumental learning. PLoS Comput. Biol. 19, (2023).

35. Rouder, J. N. & Mehrvarz, M. Random-effects psychophysics for studying individual differences in perception and cognition. Psychon. Bull. Rev. 33, (2026).

36. Engelmann, J. B. & Pessoa, L. Embedding reward signals into perception and cognition. Front. Neurosci. (2010).

37. Aston-Jones, G. & Cohen, J. D. An integrative theory of locus coeruleus-norepinephrine function: Adaptive gain and optimal performance. Annu. Rev. Neurosci. 28, 403–450 (2005).

38. Urai, A. E., Braun, A. & Donner, T. H. Pupil-linked arousal is driven by decision uncertainty and alters serial choice bias. Nat. Commun. 8, (2017).

39. Erdfelder, E., FAul, F., Buchner, A. & Lang, A. G. Statistical power analyses using G*Power 3.1: Tests for correlation and regression analyses. Behav. Res. Methods 41, 1149–1160 (2009).

40. Strasburger, H., Rentschler, I. & Jüttner, M. Peripheral vision and pattern recognition: A review. J. Vis. 11, 13 (2011).

41. Levitt, H. Transformed Up-Down Methods in Psychoacoustics. J. Acoust. Soc. Am. 49, 467–477 (1971).

42. Fong, Y., Rue, H. & Wakefield, J. Bayesian inference for generalized linear mixed models. Biostatistics 11, 397–412 (2010).

43. Geisler, W. S. Contributions of ideal observer theory to vision research. Vision Res. 51, 771–781 (2011).

44. Green, D.M & Swets, J. A. Signal detection and psychophysics. New York: Wiley (1966).

45. Kontsevich, L. L. & Tyler, C. W. Bayesian adaptive estimation of psychometric slope and threshold. Vision Res. 39, 2729–2737 (1999).

46. Aleci, C. & Rosa, C. Psychophysics in the ophthalmological practice—I. visual acuity. *Ann*. Eye Sci. 7, (2022).

47. Bishop. Pattern Recognition and Machine Learning. Pattern Recognition and Machine Learning (1971). doi:10.1007/978-1-4615-7566-5.

48. Penny, W. & Holmes, A. Random-Effects Analysis. in Human Brain Function: Second Edition 843–850 (2003). doi:10.1016/B978-012264841-0/50044-5.

49. Gilzenrat, M. S., Cohen, J. D., Rajkowski, J. & Aston-Jones, G. Pupil dynamics predict changes in task engagement mediated by locus coeruleus. Soc. Neurosci. Abstr. 515.19 (2003).

50. Palaskar & Dang. How pupil tracks cognitive processes underlying internally- and externally-directed attention tasks. in Proceedings of the Annual Meeting of the Cognitive Science Society, 46. (2024).

